# hopsy - a methods marketplace for convex polytope sampling in Python

**DOI:** 10.1101/2023.12.22.573091

**Authors:** Richard D. Paul, Johann F. Jadebeck, Anton Stratmann, Wolfgang Wiechert, Katharina Nöh

**Affiliations:** Institute of Bio- and Geosciences, IBG-1: Biotechnology, Forschungszentrum Jülich, Wilhelm-Johnen-Str., 52428 Jülich, Germany; Institute of Advanced Simulations, IAS-8: Data Analytics and Machine Learning, Forschungszentrum Jülich, Wilhelm-Johnen-Str., 52428 Jülich, Germany; Computational Systems Biotechnology (AVT.CSB), RWTH Aachen University, Forckenbeckstr., 52074 Aachen, Germany

## Abstract

**Summary:** Effective collaboration between developers of Bayesian inference methods and users is key to advance our quantitative understanding of biosystems. We here present `hopsy`, a versatile open source platform designed to provide convenient access to powerful Markov chain Monte Carlo sampling algorithms tailored to models defined on convex polytopes (CP). Based on the high-performance C++ sampling library `HOPS`, `hopsy` inherits its strengths and extends its functionalities with the accessibility of the Python programming language. A versatile plugin-mechanism enables seamless integration with domain-specific models, providing method developers with a framework for testing, benchmarking, and distributing CP samplers to approach real-world inference tasks. We showcase `hopsy` by solving common and newly composed domain-specific sampling problems, highlighting important design choices. By likening `hopsy` to a marketplace, we emphasize its role in bringing together users and developers, where users get access to state-of-the-art methods, and developers contribute their own innovative solutions for challenging domain-specific inference problems.

**Availability and Implementation:** Sources, documentation and a continuously updated list of sampling algorithms are available at https://jugit.fz-juelich.de/IBG-1/ModSim/hopsy, with Linux, Windows and MacOS binaries at https://pypi.org/project/hopsy/.

## 1 Introduction

Models are central to systems biology, acting as gateways to generate insights, making predictions, or testing hypotheses. The types of models used are diverse, ranging from statistical to physics-based. For operating models as epistemological tools, two steps are essential: exploration of the models’ capacities to represent data and estimation of model parameters from data. For both, recent years have witnessed a surge of interest in Bayesian statistics, expressing the desired information in the form of probability density functions (PDFs), under the notion of uncertainty (Wilkinson, 2006).

In many cases, the model definition spaces are (explicitly or implicitly) bounded by linear half-spaces making up a convex polytope (CP), for reasons as diverse as physiological limitations, energetic or other resource constraints, or mass balances operated at steady-state (Liebermeister and Noor, 2021). Premier examples are (bio)chemical reaction networks (Heinken et al., 2023), or ecosystem models (Gellner et al., 2023). Actually, CP-constrained models also appear in a wide range of domains outside biology, such as gravitational lensing (Lubini et al., 2013), smart power grids (Theorell and Stelling, 2022), or transport planing (Airoldi and Blocker, 2013).

CP-constrained PDFs, defined by real-world models, are high-dimensional and rarely analytically tractable. To approximate the PDFs, Markov chain Monte Carlo (MCMC) is a well-established sampling approach, with numerous variants implemented in powerful probabilistic programming tools (Carpenter et al., 2017; Oriol et al., 2023). However, such general MCMC algorithms fail to solve CP-constrained sampling problems efficiently because neglecting the CP geometry results in high rejection rates or ineffective space exploration (Jadebeck et al., 2023).

For the special case of a uniform PDF defined on CPs, plenty of algorithms have been developed (Haraldsdóttir et al., 2017; Chalkis et al., 2021), see Jadebeck et al. (2023) for a recent overview. These approaches are based on either Hit-and-Run (Bélisle et al., 1993), or interior-bound algorithms, such as Ball or Dikin Walks (Kannan et al., 1997; Kannan and Narayanan, 2012). Here, propelled by the needs in the metabolic modeling domain, various open packages have become available (Heirendt et al., 2019; Chalkis and Fisikopoulos, 2021; Jadebeck et al., 2021; Ciomek and Kadziński, 2021). This commoditization of uniform CP-constrained sampling, along with theoretical advances (Laddha and Vempala, 2021), has empowered metabolic researchers to approach increasingly high-dimensional problems within the domain (Thiele et al., 2020; Jadebeck et al., 2023), and beyond (Gellner et al., 2023).

For non-uniform CP sampling the situation is, however, quite different. Here, despite much work on “standardized” Gaussian and log-concave PDFs exists (Kook et al., 2022; Chalkis et al., 2023), the application as well as algorithmic landscapes for CP-constrained PDF sampling are scattered. Stimulated by the successes in uniform CP sampling, sampling of general non-uniform CP-constrained PDFs should be equally simple and accessible for domain experts. In turn, such a solution empowers MCMC developers to benchmark, improve and create new algorithms using real-world applications posed by domain experts.

Borrowing from the idea of a marketplace, we present the open-source Python package `hopsy`. `hopsy` is a flexible platform for general CP-constrained PDF sampling that seamlessly connects domain-specific simulation software and modern MCMC algorithms, via minimal and expressive interfaces. Specifically, `hopsy` leverages Python to allow domain experts and MCMC researchers to quickly implement and share domain-specific MCMC sampling workflows, while offering high-performance state-of-the-art implementations and support for common and innovative applications.

## 2 Approach and Implementation

By design, `hopsy` is a “batteries-included” platform to support convenient MCMC sampling of general CP-constrained PDFs, independent of the application domain. To facilitate flexibility at a low entry barrier, `hopsy` is implemented in Python and takes advantage of the C++-library for highly optimized polytope sampling `HOPS` (Jadebeck et al., 2021). Performance critical code from `HOPS` is integrated via `pybind11` (Jakob et al., 2017), while convenience functions are implemented in Python.

In the `hopsy` sampling workflow (cf. Fig. 1), a model is specified by defining the PDF on a CP-constrained support. Then a suitable sampling algorithm is selected, configured and run. For a continuously updated listing of MCMC algorithms, we refer to https://modsim.github.io/hopsy/userguide/sampling.html#proposals. After the sampling step, convergence diagnostics and visualizations are provided by the widely-used `ArviZ` package (Kumar et al., 2019).

**Fig. 1.**
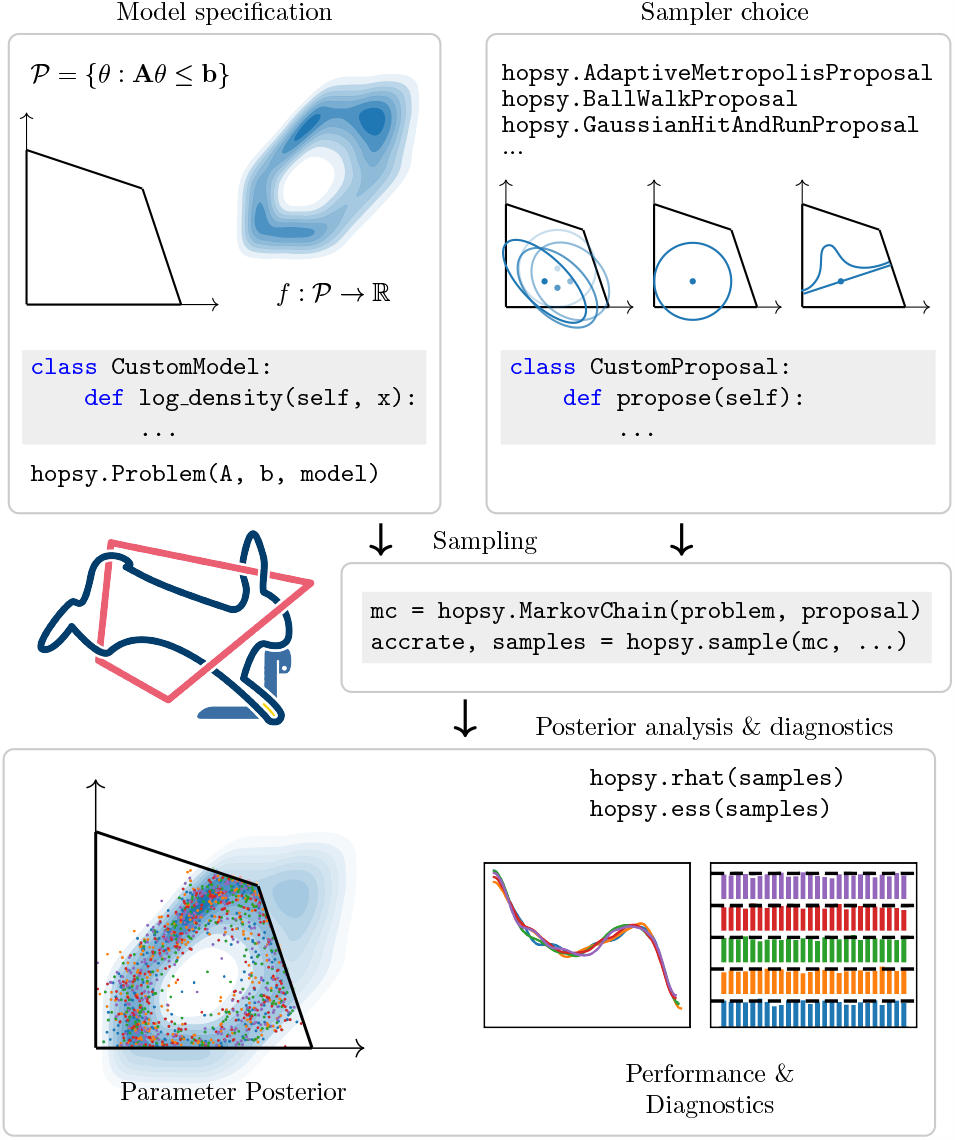
Convex polytope-constrained sampling workflow with `hopsy`.

The model specification consists of an explicit CP formulation in half-space representation and a PDF (by convention expected as log-density):

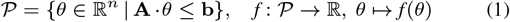

with unknown model parameters *θ* ∈ ℝ^*n*^, **A** ∈ ℝ^*n*×*m*^ and **b** ∈ ℝ^*m*^. To obtain an explicit CP formulation from an implicit one, e.g. an underdetermined linear equation system, linear algebra recipes exist (cf. SI Sec. S.2). The log-density *f* may be given as the logarithm of a closed-form PDF or, in the case of Bayesian inference, as a log-posterior provided by a simulation code. Note that the model specification in (1) is well-posed only if the resulting density function is integrable. In cases where the gradient or the curvature of the log-density *f* is available, these can be utilized for proposal construction, e.g. for Riemannian-type MCMC algorithms (Gatmiry and Vempala, 2022). Ready-to-use model specifications for standard polytopes (simplices, Birkhoff polytopes), and common log-densities are also available (cf. documentation at https://modsim.github.io/hopsy/).

The central entity that encapsulates the model specification (1) is the `hopsy`-class `hopsy.Problem`. The CP is passed in the form of domain-agnostic `NumPy` arrays (Harris et al., 2020) and the log-density is a Python object with a `log_density`-method, as well as optionally, `log_gradient` and/or positive-definite `log_curvature` methods. `hopsy`’s open plugin-architecture for custom Python code makes the formulation of the model ingredients extremely flexible. Thereby, models are either directly specified in Python or by calling external ODE- or PDE-solvers, facilitating the construction of composite models. The high flexibility for posing the log-density is achieved by a combination of interface classes and a *trampoline* (Jakob et al., 2017), which redirects calls from `HOPS` to custom log-densities defined in Python. Using the same trampoline-technique, `hopsy` supports custom MCMC implementations, which seamlessly integrate with model specifications. Importantly, this eliminates the need to write tedious “glue-code” to connect existing implementations for (1) to newly designed proposals. Examples of how easy it is to integrate and combine custom PDFs and tailor-made MCMC algorithms are given in the next section.

The model specification and user-selected sampling algorithm are collected in the `hopsy.MarkovChain` class. The function `hopsy.sample` advances a set of Markov chains by the number of requested samples. `hopsy` generates random numbers using statistically reliable 64 bit permuted congruential generators (O’Neill, 2014) with 128 bit states and periods of 2^128^, which is sufficient to prevent repeat of random numbers in realistic settings. Because instances of `hopsy.MarkovChain` support the `pickle` functionality of Python, check-pointing sampling runs is simple. The `hopsy.sample` function stores samples either in-memory as `NumPy`-arrays or in (remote) databases (Osthege and PyMC Developers, 2023).

Further highlight features of `hopsy` are CP pre-processing by rounding using `PolyRound` (Theorell et al., 2021), feasibility checks for CPs, MCMC acceptance-rate tuning (Roberts and Rosenthal, 2001), parallel tempering (PT) (Gelman, 2014) for tackling multi-modal log-densities, and Reversible Jump MCMC (Theorell and Nöh, 2019). Software quality is ensured by a continuous integration (CI) pipeline for automatic testing, automatic code-style checks using `pre-commit`, changelogs, documentation including example Jupyter notebooks, and semantic versioning.

## 3 Showcasing hopsy

To demonstrate the flexibility of `hopsy`, we selected the field of metabolic modeling, which stands out due to its high-dimensional inference tasks within a plethora of different inference approaches.

Uniform CP sampling is a common approach for unbiased exploration of metabolic capabilities (Haraldsdóttir et al., 2017) (SI Sec. S.2). Compared to the state-of-the-art Coordinate Hit-and-Run with Rounding and Thinning (CHRRT) algorithm implemented in `HOPS` (Jadebeck et al., 2023), we ponder on alternate proposals, specifically Over-relaxed Hit-and-Run (de Concini and de Martino, 2015), and a recent adaptive sampling approach that uses iterative singular value decomposition (SVD) transformations of the parameter space instead of polytope rounding (Chalkis et al., 2021). Exploiting the flexibility of the open plugin-architecture, implementing the alternate proposals took only few lines of code (30 for Over-relaxed Hit-and-Run and 49 for iterative SVD-based rounding). We benchmarked the three MCMC algorithms for a synthetic problem (16D Birkhoff polytope), and two *E. coli* models. For the synthetic problem and one of the *E. coli* models the adaptive sampling approach were the most efficient, while for the other *E. coli* model CHRRT was found best (cf. Fig. S.2). The result underlines the intricacy of CP sampling and, thus, the importance of proposal design and testing, even for similar models of the same organism.

A key quantity in metabolic models are reaction rates (fluxes) that are inferred from isotopic labeling data, in a framework known as Bayesian ^13^C-MFA (Theorell et al., 2017) (SI Sec. S.3). Here, the simulation of ^13^C-labeled metabolites entertains a nonlinear mapping from the flux parameter space to the observation space. consequently, the sum-of-squares residual between simulated and measured ^13^C-labeled metabolites gives rise to a non-uniform CP-constrained PDF. Since efficient simulation is key, the ^13^C-model was called via the domain-specific high-performance simulator `13CFLUX2` (Weitzel et al., 2012). The Hit-and-Run with Rounding algorithm was used without and with PT for sampling one of the above-mentioned *E. coli* models. Our results show that (1) PT led to improved mixing, as visible in the trace plots and Gelman-Rubin diagnostic (cf. Fig. S.4), and (2) compared to uniform sampling, incorporating ^13^C data strongly reduced the flux parameter uncertainty (cf. Fig. S.5), and predict likely measurement distributions for isotope labeling experiments (cf. Fig. S.6).

In a third application, we highlight the flexibility of `hopsy` by creating a composite model using the trampoline (SI Sec. S.4). Conventionally, extracellular rates are first estimated by means of so-called bioprocess models. Once estimated, they are incorporated into the ^13^C-model as external rate measurements together with their (symmetrized) standard deviations. From a statistical standpoint, simultaneous estimation of bioprocess and ^13^C-model parameters promises a better understanding of parameter correlations. Therefore, in a rapid prototyping manner, we built a new composite model with which bioprocess and flux parameters are estimated simultaneously. Comparing the outcome with the conventional modeling procedure in Fig. S.6 indeed shows an information gain. Thus, with `hopsy` it was simple to quickly test a new promising modeling idea in the field of ^13^C-MFA.

## 4 Conclusion

`hopsy` is an mature open source Python toolbox for highly optimized polytope sampling. `hopsy` strips the polytope sampling problem down to its minimal formal requirements and provides a simple, extensible interface. This makes `hopsy` applicable to a broad range of polytope sampling problems, such as exploring polytopic parameter spaces by uniform sampling, testing MCMC approaches, and efficiently tackling challenging Bayesian inference problems. Furthermore, we demonstrate that it is easy with `hopsy` to implement novel (composite) modeling approaches. Our showcases, despite being from the field of metabolic modeling, are inspirations for other scientific fields, such as ecological modeling (Gellner et al., 2023), optimization of chromatography pipelines (Schmölder and Kaspereit, 2020), or single-cell analysis (Paul et al., submitted), unlocking a broad portfolio of applications for method developers. `hopsy` thus fertilizes collaboration between domain experts and MCMC developers by facilitating easy sharing of new problems and MCMC approaches, allowing both communities to bring their fruits into practice quicker.

## Supporting information

Main Manuscript

## Acknowledgments

The authors thank Martin Beyß, Stephan Noack and Jochen Nießer for helpful discussions.

## References

Airoldi EM, Blocker AW. Estimating latent processes on a network from indirect measurements. JASA, 108(501):149–164, 2013.

Bélisle CJP, Romeijn HE, Smith RL. Hit-and-Run algorithms for generating multivariate distributions. Math Oper Res, 18(2):255–266, 1993.

Carpenter B et al. Stan: A probabilistic programming language. J Stat Softw, 76(1), 2017.

Chalkis A et al. Geometric algorithms for sampling the flux space of metabolic networks. In Buchin K, de Verdière ÉC, eds, 37th Int. Symp. on Computational Geometry (SoCG 2021), vol. 189 of Leibniz Int. Proc. Inform. (LIPIcs), 21:1–21:16, Dagstuhl, Germany, 2021.

Chalkis A, Fisikopoulos V. VolEsti: Volume approximation and sampling for convex polytopes in R. R J, 13(2):642–660, 2021.

Chalkis A et al. Truncated log-concave sampling for convex bodies with reflective hamiltonian monte carlo. ACM Trans Math Softw, 49(2), 2023.

Ciomek K, Kadziński M. Polyrun: A Java library for sampling from the bounded convex polytopes. SoftwareX, 13:100659, 2021.

de Concini G, de Martino D. Over-relaxed Hit-and-Run Monte Carlo for the uniform sampling of convex bodies with applications in metabolic network biophysics. Int J Mod Phys C, 26(1):1550010, 2015.

Gatmiry K, Vempala SS. Convergence of the Riemannian Langevin algorithm. ArXiv, abs/2204.10818, 2022.

Gellner G et al. Stable diverse food webs become more common when interactions are more biologically constrained. PNAS, 120(31):2017, 2023.

Gelman A. Bayesian Data Analysis. CRC Press, Boca Raton, 2014.

Haraldsdóttir HS et al. CHRR: Coordinate Hit-and-Run with rounding for uniform sampling of constraint-based models. Bioinformatics, 33(11): 1741–1743, 2017.

Harris CR et al. Array programming with NumPy. Nature, 585(7825): 357–362, 2020.

Heinken A et al. Genome-scale metabolic reconstruction of 7,302 human microorganisms for personalized medicine. Nat Biotechnol, 41(9): 1320–1331, 2023.

Heirendt L et al. Creation and analysis of biochemical constraint-based models using the COBRA toolbox v.3.0. Nat Protoc, 14(3):639–702, 2019.

Jadebeck JF et al. HOPS: High-performance library for (non-) uniform sampling of convex-constrained models. Bioinformatics, 37(12):1776–1777, 2021.

Jadebeck JF et al. Practical sampling of constraint-based models: Optimized thinning boosts chrr performance. PLOS Comput Biol, 19 (8):e1011378, 2023.

Jakob W et al. pybind11 – Seamless operability between C++11 and Python, 2017. https://github.com/pybind/pybind11.

Kannan R, Narayanan H. Random walks on polytopes and an affine interior point method for linear programming. Math Oper Res, 37(1): 1–20, 2012.

Kannan R et al. Random walks and an O*(n5) volume algorithm for convex bodies. Random Struct Algorithms, 11:1–50, 1997.

Kook Y et al. Sampling with Riemannian Hamiltonian Monte Carlo in a constrained space. In Koyejo S et al., editors, Adv Neural Inf Process Syst, 35:31684–31696. Curran Associates, Inc., 2022.

Kumar R et al. Arviz a unified library for exploratory analysis of Bayesian models in python. JOSS, 4(33):1143, 2019.

Laddha A, Vempala SS. Convergence of Gibbs sampling: Coordinate Hit-and-Run mixes fast. In Buchin K, de Verdière ÉC, eds, 37th Int. Symp. on Computational Geometry (SoCG 2021), vol. 189 of Leibniz Int. Proc. Inform. (LIPIcs), 51:1–51:12, Dagstuhl, Germany, 2021.

Liebermeister W, Noor E. Model balancing: A search for in-vivo kinetic constants and consistent metabolic states. Metabolites, 11(11):749, 2021.

Lubini M et al. Cosmological parameter determination in free-form strong gravitational lens modelling. MNRAS, 437(3):2461–2470, 2013.

O’Neill ME. PCG: A family of simple fast space-efficient statistically good algorithms for random number generation. Technical Report HMC-CS-2014-0905, Harvey Mudd College, Claremont, CA, 2014.

Oriol A-P et al. PyMC: A modern and comprehensive probabilistic programming framework in Python. PeerJ Comput Sci, 9:e1516, 2023.

Osthege M et al. McBackend, GitHub repository, https://github.com/pymc-devs/mcbackend, v0.5.0, 2023.

Paul RD et al. Robust approximate characterization of single-cell heterogeneity in microbial growth. (submitted).

Roberts GO, Rosenthal JS. Optimal scaling for various Metropolis-Hastings algorithms. Stat Sci, 16(4):351–367, 2001.

Schmölder J, Kaspereit M. A modular framework for the modelling and optimization of advanced chromatographic processes. Processes, 8(1): 65, 2020.

Theorell A, Nöh K. Reversible jump MCMC for multi-model inference in metabolic flux analysis. Bioinformatics, 36(1):232–240, 2019.

Theorell A, Stelling J. Metabolic networks, microbial consortia, and analogies to smart grids. Proc. IEEE, 110(5):541–556, 2022.

Theorell A et al. To be certain about the uncertainty: Bayesian statistics for 13C metabolic flux analysis. Biotechnol Bioeng, 114(11):2668–2684, 2017.

Theorell A et al. Polyround: polytope rounding for random sampling in metabolic networks. Bioinformatics, 38(2): 556–567, 2021.

Thiele I et al. Personalized whole-body models integrate metabolism, physiology, and the gut microbiome. Mol Syst Biol, 16(5), 2020.

Weitzel M et al. 13CFLUX2 — high-performance software suite for 13C-metabolic flux analysis. Bioinformatics, 29(1):143–145, 2012.

Wilkinson DJ Bayesian methods in bioinformatics and computational systems biology. Brief Bioinf, 8(2):109–116, 2006.

