## Supplementary material for "hopsy - a methods marketplace for convex polytope sampling in Python": Main Manuscript

#### S.1 Code availability

The `hopsy` source code is available at `https://jugit.fz-juelich.de/IBG-1/ModSim/hopsy` under the MIT license with documentation `https://modsim.github.io/hopsy/`. Available sampling algorithms are continuously updated at `https://modsim.github.io/hopsy/userguide/sampling.html\#proposals`. The code for all example workflows described here is available at `https://jugit.fz-juelich.de/IBG-1/ModSim/Fluxomics/hopsy-publication`. This includes jupyter notebooks to reproduce all figures and all model files, accessible in the `models` directory. Further source code is available in the `src` subdirectory.

#### S.2 Uniform Convex Polytope Sampling

As an example for deriving an explicit convex polytope (CP) formulation from an implicit one, we consider the case of Metabolic Flux Analysis (MFA). MFA metabolic networks consist of a set of biochemical reactions. The evolution of intracellular metabolite concentrations $X$ over time is described by a system of ordinary differential equations (ODE). These ODEs are derived from mass balancing of each reaction and parameterized by the metabolic reaction rates $\nu$, which are also referred to as *fluxes*. The stoichiometric coefficients of the metabolites in each reaction are collected in the stoichiometric matrix $S$. Consequently, the ODE system is given as

$$\frac{\mathrm{d}X}{\mathrm{d}t} = S \cdot \nu \tag{1}$$

At metabolic steady state, the intracellular metabolic concentrations are constant. In this case, the dynamic system becomes an underdetermined linear equation system for the unknown fluxes $\nu$:

$$0 = S \cdot \nu \tag{2}$$

The null space of $S$ is parameterized by a subset of *free* fluxes $\theta$, which are confined to a CP

$$\mathcal{P} = \{\theta : A \cdot \theta \leq b\} \tag{3}$$

Any $\theta \in \mathcal{P}$ is a feasible solution for Equation (2). Given no additional information, all solutions of Equation (2) are equally valid. In this case, from a probabilistic perspective, the feasible solutions are encoded by the uniform distribution over $\mathcal{P}$. In the case of Bayesian inference, the uniform distribution over *polytope* is a *proper* prior, if and only if the CP is bounded, which is generally ensured in practice. Within the specification of a `hopsy.Problem`, this corresponds to choosing a constant density $f(\theta) = \text{const.}$. This case is implicitly assumed, if no `hopsy.Model` is passed to the `hopsy.Problem`.

##### S.2.1 A Toy Example

A toy example network is depicted in Figure 1. Balancing the in- and outgoing fluxes of the internal metabolite M gives the stoichiometric matrix

$$S = \begin{array}{c} \begin{matrix} u & x & y & z \end{matrix} \\ \begin{pmatrix} 1 & -1 & -1 & -1 \end{pmatrix} M, \end{array} \tag{4}$$

Normalizing by the uptake rate $u$ and assuming the fluxes $x, y, z$ only go in the nominal direction, we get

$$1 = \hat{x} + \hat{y} + \hat{z} \quad \text{and} \quad 0 \leq \hat{x}, \hat{y}, \hat{z} \leq 1 \tag{5}$$

as the solution of Equation (2) with $\hat{x} = x/u, \hat{y} = y/u, \hat{z} = z/u$. Selecting $\hat{x}$ and $\hat{y}$ as free parameters, the space of flux solutions is given by the inequality

$$1 \geq \hat{x} + \hat{y}, \tag{6}$$

which results in the polytope $\mathcal{P}$ being a two-dimensional simplex

$$A \cdot \theta = \begin{pmatrix} 1 & 1 \\ -1 & 0 \\ 0 & -1 \end{pmatrix} \cdot \begin{pmatrix} \hat{x} \\ \hat{y} \end{pmatrix} \leq \begin{pmatrix} 1 \\ 0 \\ 0 \end{pmatrix} = b. \tag{7}$$

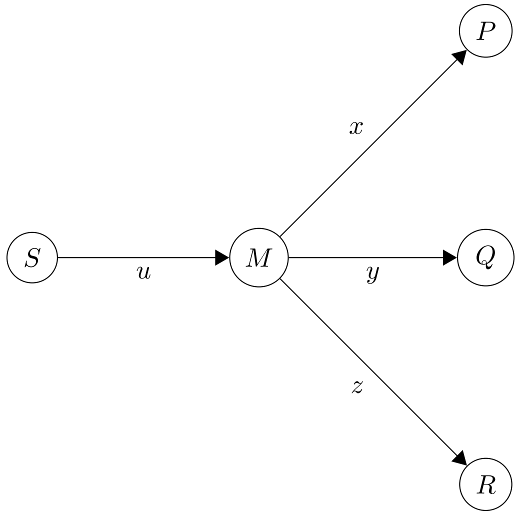

Figure 1: A simple metabolic network. Upper-case letters denote metabolites and lower-case letters are metabolic reactions. Nominal reaction directions are indicated by the arrowhead. Here, the metabolite M is considered "intracellular" and therefore mass-balanced, with a constant concentration $M$, whereas the pools S, P, Q, and R are considered extracellular metabolites with time-dependent concentrations $S, P, Q, R$.

##### S.2.2 Numerical Experiment

We used the problem of feasible flux space sampling to highlight `hopsy`'s capabilities to rapidly prototype and implement MCMC algorithms. In particular, we implemented a updated version of the little-considered Over-relaxed Hit-&-Run algorithm by [3] and a very recent iterative MCMC algorithm [2]:

- Over-relaxed Hit-&-Run with Rounding (ORHR),
- Multiphase Monte-Carlo algorithm using a Billiard walk proposal (MMBW)

We benchmarked both algorithms alongside `hopsy`'s native Coordinate Hit-&-Run with Rounding and Thinning algorithm (CHRRT) by uniformly sample three CP problems:

- a 16-dimensional Birkhoff polytope (`BP5`)
- a 18-dimensional $^{13}C$ metabolic flux model of *E. coli* from [15] (`zamboni`)
- a 24-dimensional constraint-based *E. coli* model from [9] (`e_coli_core`)

See `https://gitlab-public.fz-juelich.de/IBG-1/ModSim/Fluxomics/hopsy-publication/-/blob/main/Eval-use-case-1.ipynb` for definitions of the polytopes and further details regarding this benchmark.

All calculations were run on a system equipped with AMD 5950x CPU and running Ubuntu 22.04. For every of the three sampling algorithms, we ran four parallel chains, parallelized using Python's `multiprocessing` library. For the comparison of the three algorithms, each chain was run until an effective sample size (ESS) [4] of at least 1,000 was achieved. According to [6], the selection of the thinning factor impacts the algorithm performance considerably when rounding is involved, primarily due to the de-rounding step that maps the set of thinned samples back to the original parameter space. Here, following the guideline derived in [6], the thinning factor was set to $\textit{problem\_dimension}^2 \times 1./6$ for CHRRT and ORHR. Because the MMBW algorithm rounding works differently, thinning was not applied in this case.

##### S.2.3 Results

Figure 2 shows the relative ESS per sample and per unit of time (in seconds) as measures of efficiency of three sampling algorithms used. Considering both measures, the Multiphase Billiard algorithm MMBW has the highest performance for `BP5` and `e_coli_core`. However, for the `zamboni` model, the CHRRT algorithm,

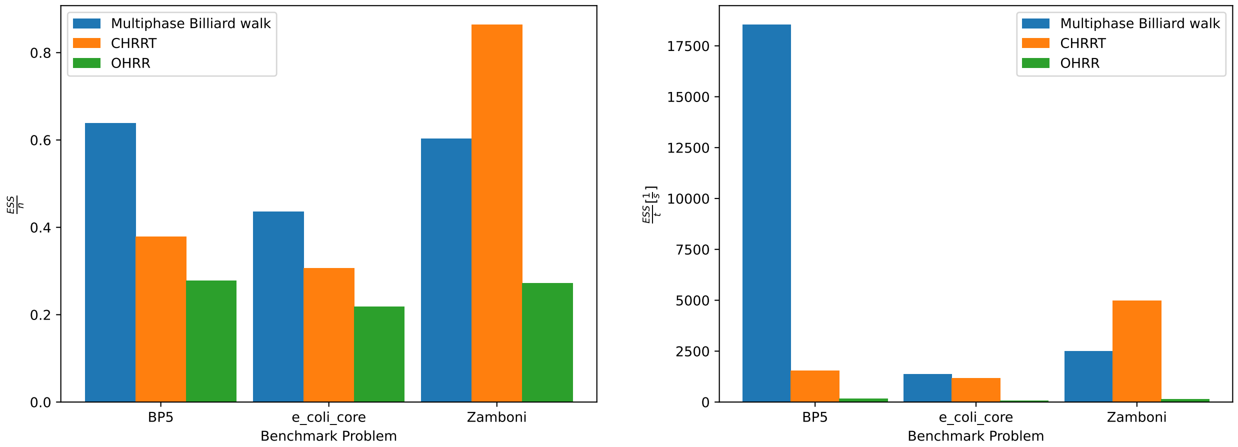

Figure 2: Benchmark results. Relative effective sample size (ESS) per sample (left) and time measured in seconds (right) for the three algorithms (MMBW, CHRRT, ORHR) run on three problems (`BP5`, `e_coli_core`, `zamboni`).

which is built-in in `hopsy`, is most efficient. The marginal flux distributions of the three uniform sampling problems are shown in Figure 3. Note that due to the geometry of the CPs, the marginal distributions are seldom uniform.

Both `e_coli_core` and `zamboni` are core models of *E. coli*, yet different algorithms perform best, showing that the performance of MCMC algorithms is problem specific. This motivates the development and inclusion of new algorithms in `hopsy`. Indeed, in view of the results in Figure 2, the MMBW algorithm was natively included in `hopsy` version 1.4.0.

(a) 16-dimensional Birkhoff polytope BP5.

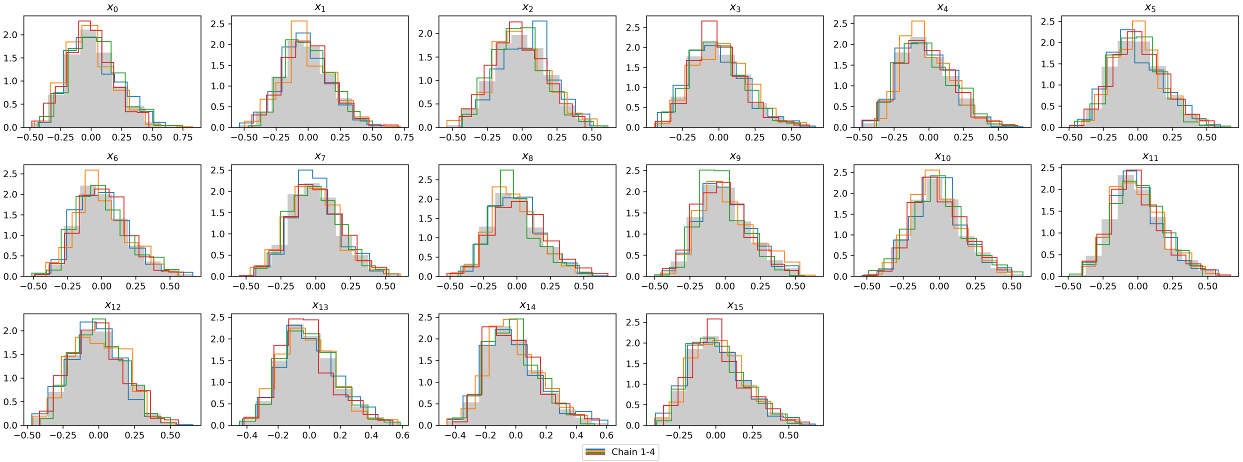

(b) *E. coli* model zamboni.

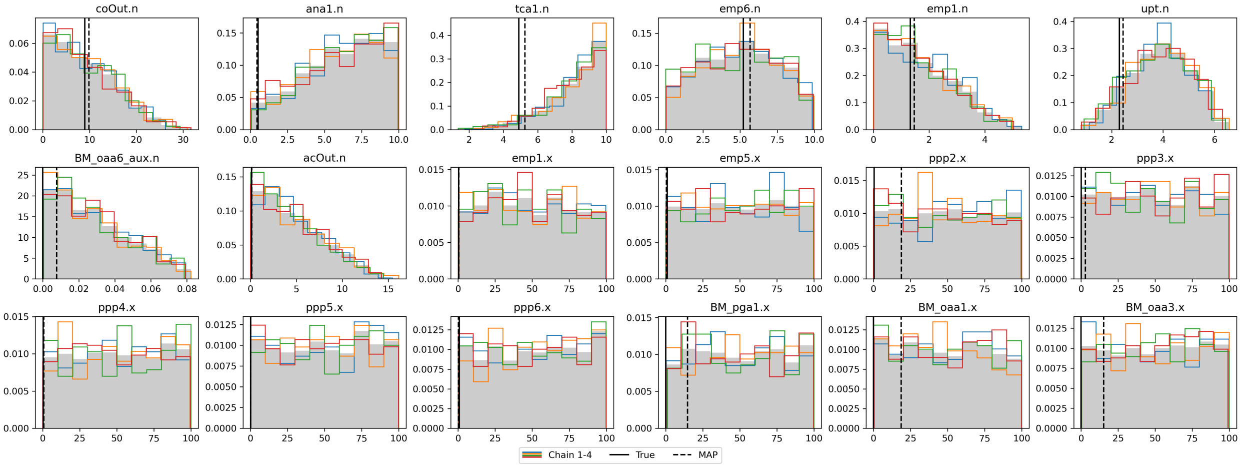

(c) *E. coli* model e_coli_core

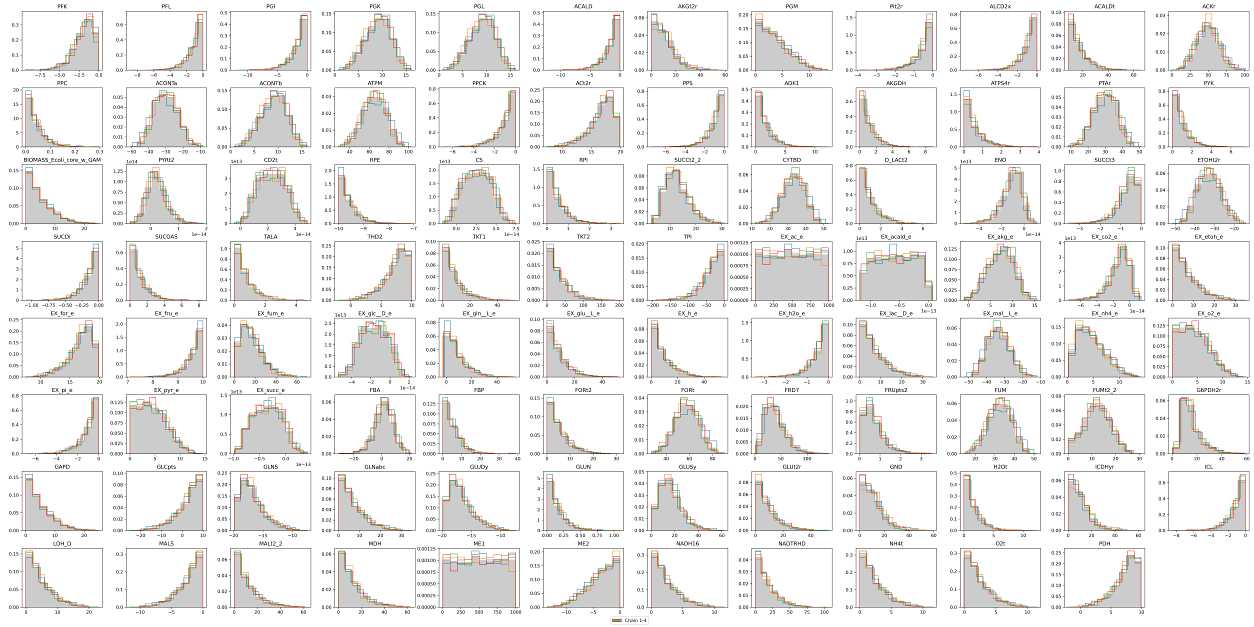

Figure 3: Marginal distributions of random variables uniformly distributed on CP test problems. In (c) the full flux space, consisting of 95 fluxes, is shown.

#### S.3 Bayesian $^{13}$C-Metabolic Flux Analysis

Uniform flux space sampling, as described in Sec. S.2, represents the constraining impact of the model formulation on the fluxes, without considering measurements. In Bayesian $^{13}$C-Metabolic Flux Analysis (MFA), two data types enter the inference problem. One data type is provided by the so-called *extracellular rates*. These rates represent the biomass-specific substrate uptake and product excretion rates, as well as the growth rate of the organism under study. The extracellular rates are extracted from exometabolomic concentration data, either by metabolite-wise linear regression [8] or by consistent inference using bioprocess models [5] (see also Sec. S.4).

While the knowledge of the extracellular rates is helpful, they are insufficient for determining the *intracellular fluxes* in parallel and cyclic metabolic pathways [14]. For this, the second data type is $^{13}$C labeling data of intracellular metabolites that are acquired by administering carbon-based substrates in isotope labeling experiments. Specifically, $^{13}$C-labeled substances are taken up by the cells, the label is propagated through the metabolic pathways according to the underlying fluxes, and chemical analytics (mass spectrometry or NMR) allows for the quantification of fractional enrichments of internal metabolites. On the other hand, the metabolic model is complemented with a carbon atom transition network, describing the fate of the individual carbon atoms through the metabolic network. This model is used to simulate the measurable isotope labeling states. Together, this allows estimating the fluxes by means of a computational procedure. Bayesian $^{13}$C-MFA uses Bayes theorem to produce flux posterior probability distributions, see [11] for details.

The measurement errors on extracellular rate and labeling data are both modelled to be additive Gaussian. Formally, let $Y_{13C}$ be the data vector of extracellular rate and isotope enrichment measurements, then

$$Y_{13C} \sim \mathcal{N}(g(\theta), \Sigma_{13C}) \tag{8}$$

where $g(\theta)$ is the vector of computed extracellular rates and simulated isotope labeling, given known fluxes $\theta$, and $\Sigma_{13C}$ is the associated diagonal measurement covariance matrix. Note, that the definition space of $\theta$ is restricted to a CP as detailed in Sec. S.2. Using the uniform distribution on the flux CP as prior, the log-density function for Bayesian $^{13}$C-MFA (up to a constant) then is

$$\log f(\theta) = \frac{1}{2} \|Y_{13C} - g(\theta)\|^2_{\Sigma_{13C}} \tag{9}$$

Since simulation of isotope data by evaluating $g(\theta)$ is computationally expensive, we rely on the high-performance simulator 13CFLUX2 [13] for fast log-density evaluation.

##### S.3.1 Numerical Experiment

Using the `zamboni` model, we simulated isotope labeling data for an input tracer of [1,2]-$^{13}$C-labeled glucose. The reference fluxes and the measurement configuration were taken from [15] to simulate ground truth data, see model file at `https://jugit.fz-juelich.de/IBG-1/ModSim/Fluxomics/hopsy-publication/-/blob/main/models/Ecoli_NatProt_perturbed_small.fml?ref_type=heads`. The simulated rate and isotope labeling measurements were perturbed according to the Gaussian error model in Equation (8) using the standard deviation reported in [15].

We used an Intel(R) Xeon(R) CPU E5-2683 v4 @ 2.10GHz running Ubuntu 22.04 to simulate 64 chains in parallel twice. In the first (non-PT) run, all chains were independent. In the second (PT) run, we let the chains communicate by binding the chains together in sets of 16. In both experiments, we sampled 1e6 samples with a thinning of 20 using the Hit-and-Run with Rounding algorithm and discarded the first half of the chains as burn-in.

##### S.3.2 Results

Both MCMC runs, PT and non-PT, converged, as indicated by the potential scale reduction factor $\hat{R}$ [4], and the trace plots in Figure 4. For the independent chains, the $\hat{R}$ was below 1.02 after 10.8 h, indicating convergence. The PT run achieved an $\hat{R}$ below 1.002 in 12.8 h, therefore mixed better for the same number of samples drawn with around 1.2 × longer run-time.

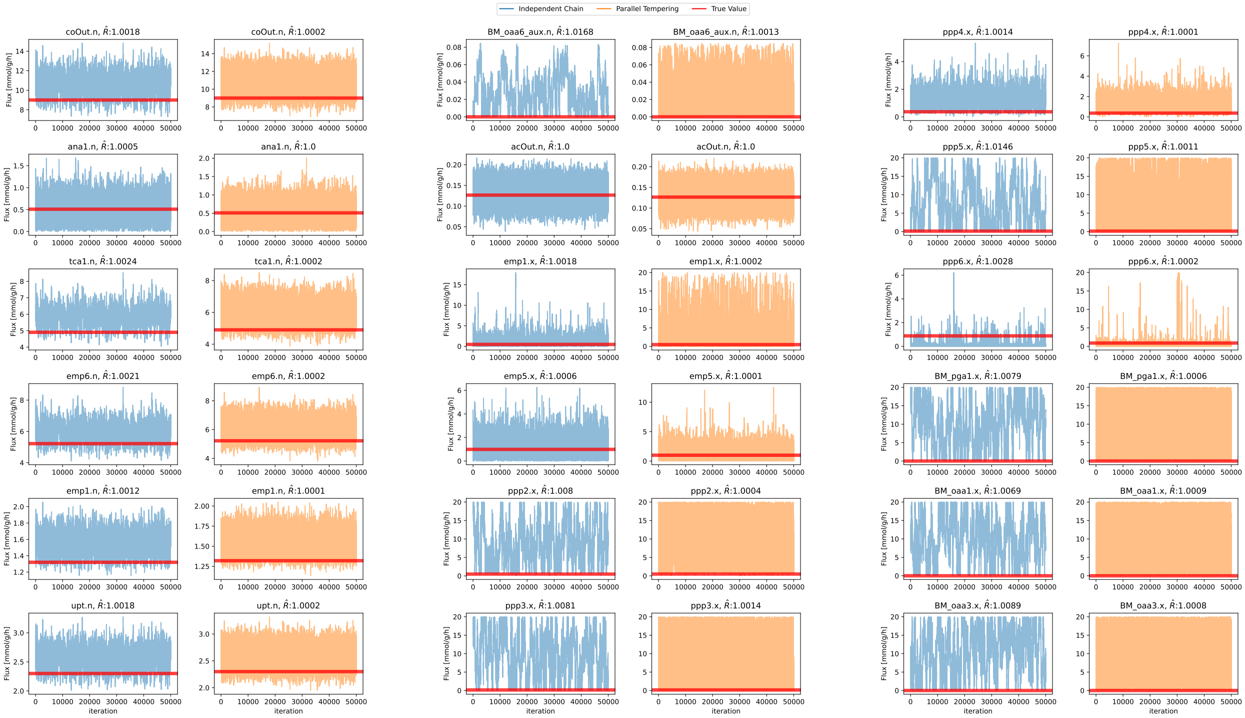

Figure 4: Trace plots of MCMC simulations without (blue) and with parallel tempering (orange). The horizontal red line represents the reference fluxes. High confidence in MCMC convergence is visible as a 'lack of structure" in the trace plots, i.e. little auto-correlation of the fluxes. For some fluxes, such as `ppp2.x`, the reduction in auto-correlation from using parallel tempering is clearly visible. For each flux, the rank normalized potential scale reduction factors were computed using `Arviz` [7].

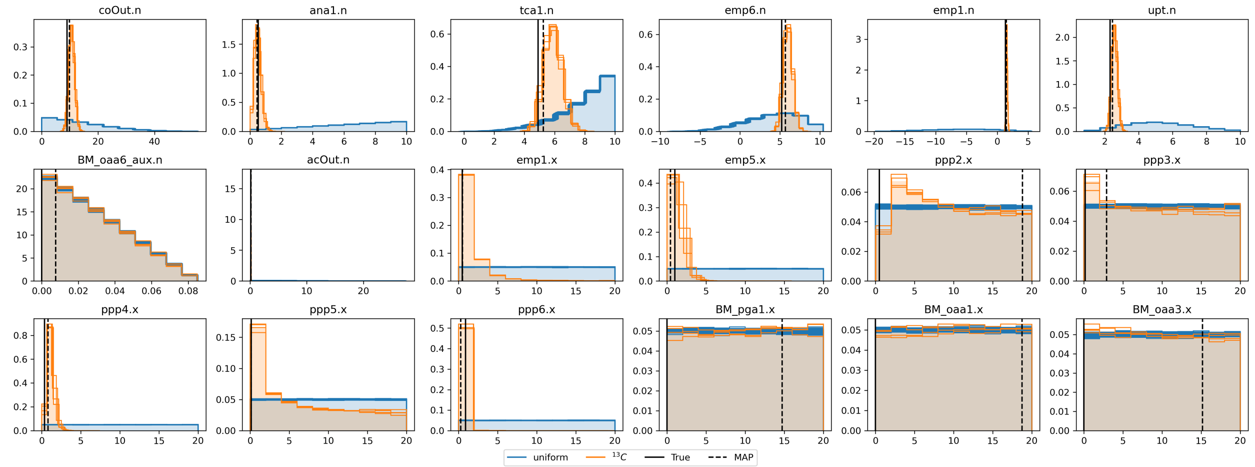

Figure 5: Comparison of marginal distributions for the independent fluxes of the `zamboni` model. Uniform (prior) marginal distributions (blue) and posterior marginal distributions (orange). The reference fluxes are indicated by a vertical dashed line. The comparison emphasizes the information gain in major fluxes (primarily net fluxes) obtained from taking $^{13}$C-MFA data into account.

The marginal distributions obtained from sampling the $^{13}$C-MFA posterior (Equation (9)) are shown in Figure 5. To study the information gain obtained from taking extracellular rates and isotope labeling data into account, we compare the marginal flux posterior distributions with the marginal flux prior distributions (cf. Section S.2). Except for six of the 18 fluxes, one net flux (`BM_oaa6_aux.n`) and five exchange fluxes (`ppp2.x`, `ppp3.x`, `BM_pga1.x`, `BM_oaa1.x`, `BM_oaa3.x`), a clear reduction in variance is observable. Notice that exchange fluxes are indeed notoriously hard to identify [14].

Finally, we show how sampling results contribute to experimental design decisions. As an example, Figure 6 shows the isotope labeling data predicted for [1,2]-$^{13}$C-labeled glucose as tracer, by re-using the results generated by uniform sampling before (cf. Sec. S.2). In the same way, arbitrary tracer mixture can be tested, predicting likely isotope labeling patterns to emerge, being a valuable information source for analytical experts [10].

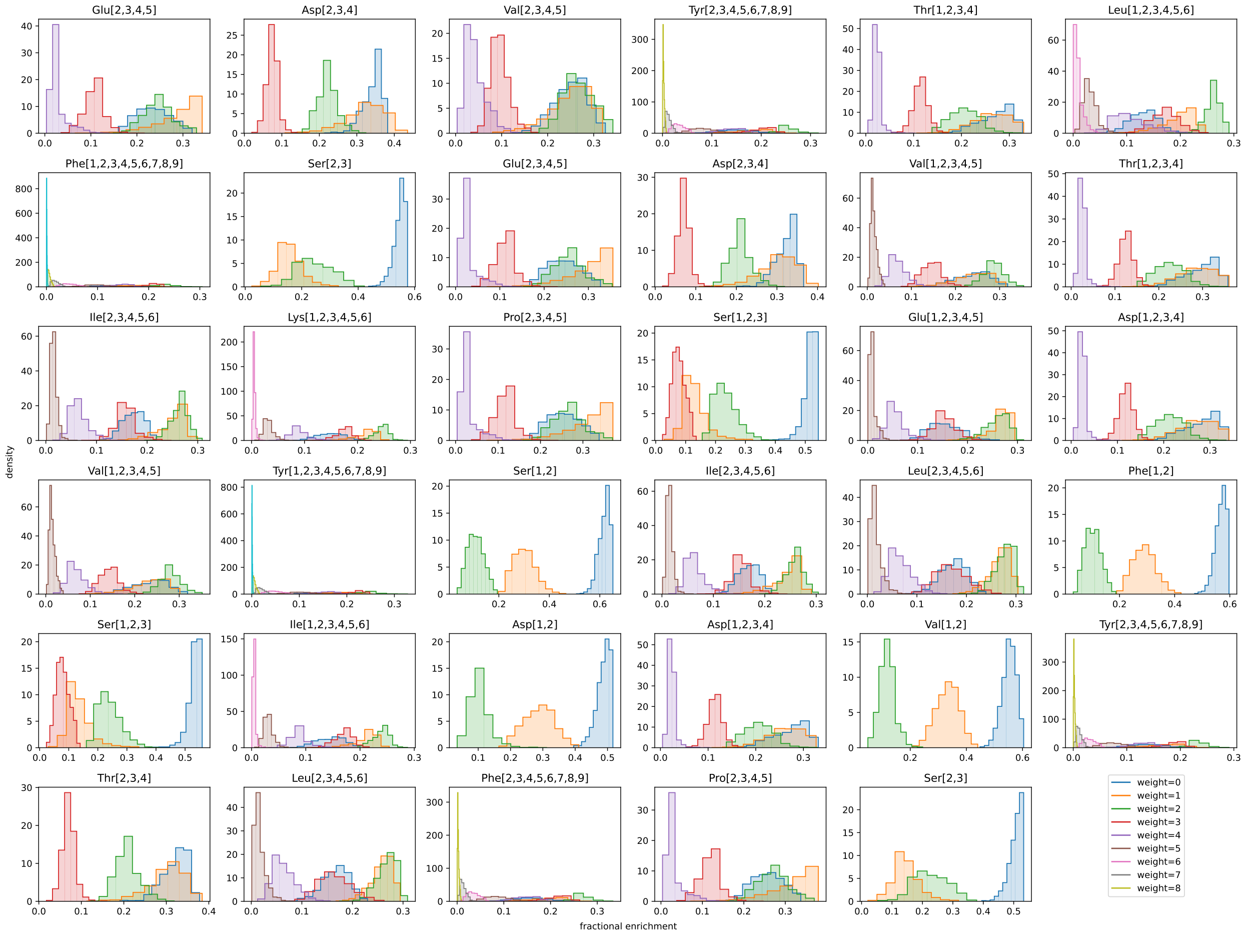

Figure 6: Sample-based prediction of measurements as part of experimental design.

#### S.4 Composite Bioprocess and Metabolic Flux Modeling

In $^{13}$C-MFA, intracellular fluxes are traditionally determined sequentially: first, extracellular rates are estimated using linear regression or a bioprocess model, then, the inferred rates along with isotope labeling data are used for estimating the intracellular flux using a $^{13}$C-MFA model (cf. Sec. S.3). This sequential process, however, neglects the correlation between those parameters that are shared between the bioprocess and the $^{13}$C-MFA models, i.e. the extracellular rates. In turn, the neglection might lead to an information bias. To improve the reliability of the parameter estimates, we here, for the first time, simultaneously inferred bioprocess and $^{13}$C-MFA parameters, exploiting `hopsy`'s trampoline mechanism. Finally, we compared the inferences using the composite model with those of the sequential approach.

A bioprocess model is a kinetic model that balances the dynamic substance concentrations $X$ in a bioreactor, starting from initial concentrations $X_0$:

$$\frac{\mathrm{d}X}{\mathrm{d}t} = h(X, \theta_{bp}), \quad X(0) = X_0 \tag{10}$$

where $h$ is a nonlinear function of kinetic parameters $\theta_{bp}$, which describes the time-dependent "mechanistics" of biomass and extracellular compound concentrations in a cultivation, such as exponential biomass growth or Monod substrate uptake kinetics. Given a time-series of extracellular biomass and concentration measurements $Y_{bp}$ with additive Gaussian errors, we have

$$Y_{bp} \sim \mathcal{N}(X(\theta_{bp}), \Sigma_{bp}) \tag{11}$$

with $X(\theta_{bp})$ being the solution of Equation (10) for given kinetic parameters $\theta_{bp}$ and the (diagonal) measurement covariance matrix $\Sigma_{bp}$.

In contrast to bioprocess models that balance time-dependent extracellular concentrations within a bioreactor, $^{13}$C-MFA models balance the metabolite concentrations within a cell that is in a metabolic (pseudo)steady state (cf. Sec. S.3). The precondition to connect bioprocess and $^{13}$C-MFA models is, thus, that in the time interval in which isotope labeling is administered, the rate of change in extracellular concentrations $h(X, \theta_{bp})$ is approximately constant, suggesting that the cells' uptake, secretion, and growth rates remain reasonably constant. We check this criterion by computing the maximal deviation from the mean rates during the labeling interval. If the maximal deviation does not exceed a tolerable threshold, we expect that the bioprocess shows constant growth, and thus, the cells are in a metabolic steady state.

If this precondition is fulfilled, $^{13}$C-MFA (Equation (8)) and bioprocess (Equation (11)) measurements are combined to the joint measurement vector $Y$ and the joint measurement covariance matrix, resulting in the composite statistical model

$$Y = \begin{pmatrix} Y_{13C} \\ Y_{bp} \end{pmatrix} \sim \mathcal{N}\left( \begin{pmatrix} g(\theta_{13C}) \\ X(\theta_{bp}) \end{pmatrix}, \begin{pmatrix} \Sigma_{13C} & 0 \\ 0 & \Sigma_{bp} \end{pmatrix} \right) \tag{12}$$

With the joint parameter vector $\theta$ now constituted by the parameters of the bioprocess and $^{13}$C-MFA models, this gives the composite posterior log density

$$\log f(\theta) = \begin{cases} \frac{1}{2}\|Y_{bp} - X(\theta_{bp})\|^2_{\Sigma_{bp}} + \frac{1}{2}\|Y_{13C} - g(\theta_{13C})\|^2_{\Sigma_{13C}} & \text{if } h(X, \theta_{bp}) \approx \text{const. and } \theta_{13C} \in \mathcal{P}, \\ -\infty & \text{else,} \end{cases} \tag{13}$$

where $g(\theta_{13C})$ is the forward simulation of rate and labeling measurements of an isotope labeling experiment with the $^{13}$C-MFA model, as described in Section S.3.

Note, that the composite model introduces an implicit dependency between the parameters of the $^{13}$C-MFA model and the parameters of the bioprocess model, namely $\theta_{13C} = \theta_{13C}(\theta_{bp})$. That is, extracellular fluxes are computed from the bioprocess parameters, whereas intracellular fluxes remain free parameters that are to be inferred directly. We make this more explicit for actual models in Equation (17). Here, $\theta_{13C}$ has to satisfy the linear constraints introduced in Section S.2. Since the dependency is nonlinear in $\theta_{bp}$, the condition $\theta_{13C} \in \mathcal{P}$ has to be tested for every new parameter estimate $\theta_{bp}$.

##### S.4.1 Numerical Experiment

We combined the previously used `zamboni` model of *E. coli* with a chemostat bioprocess model, which is given by the following system of ordinary differential equations (adapted from [1]),

$$\frac{\mathrm{d}}{\mathrm{d}t}\begin{pmatrix} S \\ X \\ P \\ Q \end{pmatrix} = \begin{pmatrix} -\left(\frac{\mu}{Y_{XS}} + \frac{q_p}{Y_{PS}} + \frac{q_q}{Y_{QS}}\right) \cdot X + (S_f - S) \cdot D, \\ q_p \cdot X - D \cdot P, \\ (\mu - D) \cdot X, \\ q_q \cdot X - D \cdot Q, \end{pmatrix} \tag{14}$$

where $S$ is the substrate (glucose), $X$ the biomass, and $P$ (acetate) as well as $Q$ ($CO_2$) the extracellular product concentrations. Initial concentrations are $S(0) = 5000$ [mM], $X(0) = 2.5$ [g/L], $P(0) = 0.0$ [mM], and $Q(0) = 0.0$ [mM]. The substrate feed was fixed to $S_f = 10$ [mM] and the dilution rate was set equal to the growth rate, $D = \mu = 0.32$ [1/h].

Hence, the parameters $\theta_{bp}$ of the bioprocess model are the growth rate $\mu$ [1/h], yield coefficients $Y_{XS}$ [gCDW/mmol], $Y_{PS}$ [gCDW/mmol], and $Y_{QS}$ [gCDW/mmol], as well as product formation rates $q_p$ [mmol/gCDW/h] and $q_q$ [mmol/gCDW/h]:

$$\theta_{bp} = (\mu, Y_{XS}, Y_{PS}, Y_{QS}, q_q, q_p). \tag{15}$$

On the other hand, the $^{13}$C-MFA model is parameterized with the same free fluxes as selected before in Section S.3, namely

$$\begin{aligned} \theta_{13C} = ( \quad & upt.n, emp1.n, emp6.n, tca1.n, ana1.n, acOut.n, coOut.n, BM\_oaa6\_aux.n, \\ & emp1.x, emp5.x, ppp2.x, ppp3.x, ppp4.x, ppp5.x, ppp6.x, \\ & BM\_pga1.x, BM\_oaa1.x, BM\_oaa3.x) \end{aligned} \tag{16}$$

To combine the bioprocess and the $^{13}$C-MFA models, we associate extracellular substrate decline and product formation rates with the specific uptake rate, `upt.n`, and the two specific secretion net fluxes, `acOut.n` and `coOut.n`, of the $^{13}$C-MFA model. That is, we map

$$\begin{aligned} upt.n(\theta_{bp}) &= \frac{\mu}{Y_{XS}} + \frac{q_p}{Y_{PS}} + \frac{q_q}{Y_{QS}} \\ acOut.n(\theta_{bp}) &= q_p \\ coOut.n(\theta_{bp}) &= q_q, \end{aligned} \tag{17}$$

yielding the joint parameter vector

$$\begin{aligned} \theta = ( \quad & \mu, Y_{CS}, Y_{PS}, Y_{QS}, q_q, q_p, \\ & emp1.n, emp6.n, tca1.n, ana1.n, BM\_oaa6\_aux.n, \\ & emp1.x, emp5.x, ppp2.x, ppp3.x, ppp4.x, ppp5.x, ppp6.x, \\ & BM\_pga1.x, BM\_oaa1.x, BM\_oaa3.x) \end{aligned} \tag{18}$$

As the rates of the chemostat model are constant with respect to the concentration changes, constancy in growth conditions is ascertained, rendering the test in (13) naturally fulfilled.

Labeling measurements and extracellular rate measurements for the $^{13}$C-MFA model were simulated as described before in Section S.3. Bioprocess concentrations were simulated with parameters $\theta_{bp}$ that reproduces the previous $^{13}$C-MFA parameters using Equation (17). The simulated values were then perturbed with normally distributed $\epsilon \sim \mathcal{N}(0, \sigma)$, where $\sigma$ was set to 0.1 for $X$, $P$ and $Q$, 0.02 for $S$, and the $^{13}$C-MFA measurements as described in Section S.3. All bioprocess parameters were bounded within a range of $[0, 20]$. $^{13}$C-MFA model bounds were as before in Section S.2 and Section S.3. The bioprocess model Equation (10) is solved by a `SciPy` [12] integrator, the `zamboni` model is simulated with 13CFLUX2. Glue code to implement the composite model was written in Python and can be found at `https://jugit.fz-juelich.de/IBG-1/ModSim/Fluxomics/hopsy-publication/-/blob/main/src/bioprocciso_pt.py?ref_type=heads`.

We used an Intel(R) Xeon(R) Gold 6130 CPU @ 2.10GHz running Ubuntu 22.04 to simulate 64 chains in parallel. Because parallel tempering performed well for inference of the $^{13}$C-MFA model, we also use parallel tempering and let chains communicate in sets of 16, obtaining 4 replicates. We generated 1e6 samples with a thinning of 20 using the Hit-and-Run algorithm and discarded the first half of the chains as burn-in. The run converged with a potential scale reduction factor ($\hat{R}$) below 1.08 in around 55 hours. Since our case study was focused on rapid prototyping, less emphasis was laid on run time performance. Clearly, the run time can be improved by optimizing the simulation code.

##### S.4.2 Results

In Figure 7, we depict marginal distributions of fluxes obtained from sampling Equation (13). Note, that the fluxes `upt.n`, `acOut.n` and `coOut.n` are functions of the sampled bioprocess parameters $\theta_{bp}$ (cf. Equation (17)). Thus, for these fluxes we depict the marginal forward distributions obtained from applying the map Equation (17) to the posterior samples of the bioprocess parameters. Notably, we observe slightly improved identifiability for some fluxes, in particular the rates `coOut.n` and `upt.n`, but also the intracellular fluxes `tca1.n`, `emp6.n` and `emp1.n`. These results suggest that combining models as proposed here can improve flux estimations.

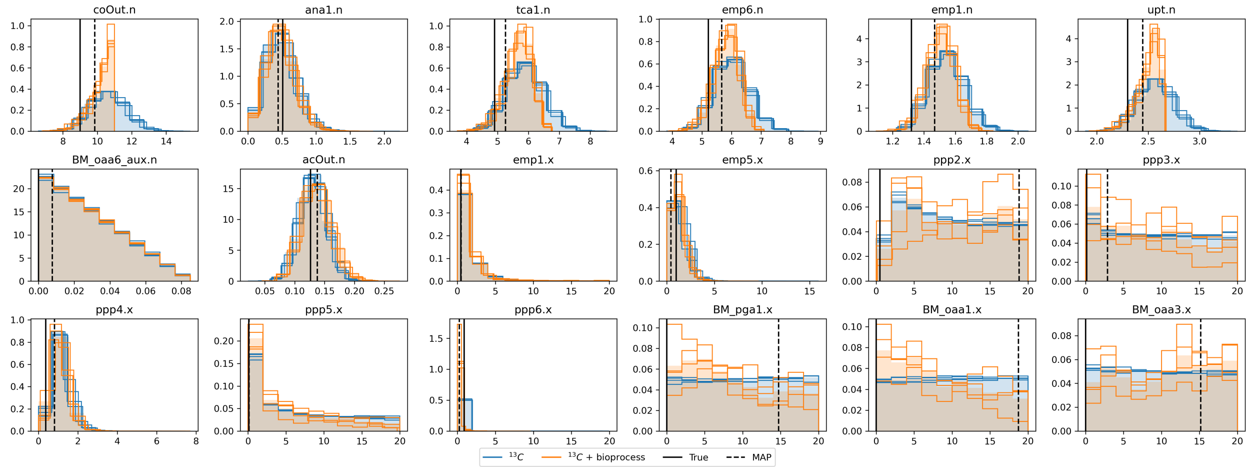

Figure 7: Comparison of posterior marginal distributions of fluxes of the `zamboni` $^{13}$C-MFA model obtained using the sequential approach, described in Section S.3, versus marginal distributions obtained from the composite modeling approach (cf. Section S.4). The black solid line indicates the true value used to generate the simulated data, the black dashed line indicates the *maximum a posteriori* (MAP) estimate of the composite model. The comparison emphasizes additional information gain in major fluxes (primarily net fluxes), obtained from taking the coupling of bioprocess and $^{13}$C-MFA modeling into account.

### References

[1] A. Ajbar and A. Fakeeha. Static and dynamic behavior of a class of unstructured models of continuous bioreactors with growth associated product. *Bioprocess and Biosystems Engineering*, 25(1):21–27, 2002.

[2] Apostolos Chalkis and Vissarion Fisikopoulos. VolEsti: Volume approximation and sampling for convex polytopes in R. *The R Journal*, 13(2):642–660, 2021.

[3] G. de Concini and D. de Martino. Over-relaxed Hit-and-Run Monte Carlo for the uniform sampling of convex bodies with applications in metabolic network biophysics. *International Journal of Modern Physics C*, 26(1):1550010, 2015.

[4] Andrew Gelman. *Bayesian Data Analysis*. CRC Press, Boca Raton, 2014.

[5] Johannes Hemmerich, Niklas Tenhaef, Wolfgang Wiechert, and Stephan Noack. pyFOOMB: Python framework for object oriented modeling of bioprocesses. *Engineering in Life Sciences*, 21(3-4):242–257, 2021.

[6] Johann F. Jadebeck, Wolfgang Wiechert, and Katharina Nöh. Practical sampling of constraint-based models: Optimized thinning boosts CHRR performance. *PLOS Computational Biology*, 19(8):e1011378, 2023.

[7] Ravin Kumar, Colin Carroll, Ari Hartikainen, and Osvaldo Martin. Arviz a unified library for exploratory analysis of Bayesian models in Python. *Journal of Open Source Software*, 4(33):1143, 2019.

[8] Taylor A. Murphy and Jamey D. Young. ETA: Robust software for determination of cell specific rates from extracellular time courses. *Biotechnology and Bioengineering*, 110(6):1748–1758, 2013.

[9] Jeffrey D. Orth et al. Reconstruction and use of microbial metabolic networks: the core *Escherichia coli* metabolic model as an educational guide. *EcoSal Plus*, 4(1), 2010.

[10] Jan Schellenberger et al. Predicting outcomes of steady-state $^{13}$C isotope tracing experiments using monte carlo sampling. *BMC Systems Biology*, 6(1):9, Jan 2012.

[11] Axel Theorell, Samuel Leweke, Wolfgang Wiechert, and Katharina Nöh. To be certain about the uncertainty: Bayesian statistics for $^{13}$C metabolic flux analysis. *Biotechnology and Bioengineering*, 114(11):2668–2684, 2017.

[12] Pauli Virtanen, Ralf Gommers, Travis E. Oliphant, Matt Haberland, Tyler Reddy, David Cournapeau, Evgeni Burovski, Pearu Peterson, Warren Weckesser, Jonathan Bright, Stéfan J. van der Walt, Matthew Brett, Joshua Wilson, K. Jarrod Millman, Nikolay Mayorov, Andrew R. J. Nelson, Eric Jones, Robert Kern, Eric Larson, C J Carey, İlhan Polat, Yu Feng, Eric W. Moore, Jake VanderPlas, Denis Laxalde, Josef Perktold, Robert Cimrman, Ian Henriksen, E. A. Quintero, Charles R. Harris, Anne M. Archibald, Antônio H. Ribeiro, Fabian Pedregosa, Paul van Mulbregt, and SciPy 1.0 Contributors. SciPy 1.0: Fundamental algorithms for scientific computing in Python. *Nature Methods*, 17:261–272, 2020.

[13] Michael Weitzel, Katharina Nöh, Tolga Dalman, Sebastian Niedenführ, Birgit Stute, and Wolfgang Wiechert. 13CFLUX2 — high-performance software suite for $^{13}$C-metabolic flux analysis. *Bioinformatics*, 29(1):143–145, 2012.

[14] Wolfgang Wiechert, Sebastian Niedenführ, and Katharina Nöh. A primer to $^{13}$C metabolic flux analysis. In *Fundamental Bioengineering*, pages 97–142. Wiley-VCH Verlag GmbH & Co. KGaA, 2015.

[15] Nicola Zamboni, Sarah-Maria Fendt, Martin Rühl, and Uwe Sauer. $^{13}$C-based metabolic flux analysis. *Nature Protocols*, 4(6):878–892, 2009.
